# Immunodominance Hierarchy of Endogenous BBN963 Bladder Cancer Antigens Remains Stable Under Anti-PD1 and Anti-CTLA4 Immunotherapy

**DOI:** 10.64898/2026.05.20.726664

**Authors:** Misha S. Fini, Jessica R. Alley, Steven P. Vensko, Dhuvarakesh Karthikeyan, Jin Seok Lee, Jason A. Garness, Emily Paul, Alex M. Jaeger, Nilu Goonetilleke, William Y. Kim, Benjamin G. Vincent

**Affiliations:** Department of Microbiology and Immunology, UNC Chapel Hill School of Medicine, Chapel Hill, NC, USA, 27514; Lineberger Comprehensive Cancer Center, UNC Chapel Hill School of Medicine, Chapel Hill, NC, USA, 27514; Department of Bioinformatics and Computational Biology, UNC Chapel Hill School of Medicine, Chapel Hill, NC, USA, 27514; Department of Genetics, UNC Chapel Hill School of Medicine, Chapel Hill, NC, USA, 27514; Department of Pharmacology, UNC Chapel Hill School of Medicine, Chapel Hill, NC, USA, 27514; Department of Medicine, UNC Chapel Hill School of Medicine, Chapel Hill, NC, USA, 27514; Department of Molecular Oncology, Moffitt Cancer Center, Tampa, FL, USA, 33612

## Abstract

Immune checkpoint inhibition (ICI) is clinically active against multiple cancers, including urothelial cancer at the non-muscle invasive, muscle-invasive, and metastatic stages. Despite this, large numbers of patients experience disease progression and relapse after treatment with ICI-containing regimens. Tumor antigen-specific T cells are critical to ICI response, however few studies have evaluated the breadth and magnitude of tumor antigen-specific T cell responses with ICI therapy. In this study, we mapped the tumor antigen immunodominance hierarchy in the BBN963 model of murine basal-like bladder cancer for endogenous tumor neoantigens expressed physiologically. We used a high-throughput matrixed ELISpot assay to detect CD8^+^ T cell responses to predicted BBN963 tumor antigens derived from multiple mutational genomic sources. We found CD8+ T cell responses were directed against a subset of tumor antigens forming a stable and reproducible immunodominance hierarchy across individual mice. Treatment with anti-PD-1 or anti-CTLA-4 did not substantially reshape this hierarchy or broadly shift dominant responses to previously defined subdominant epitopes. Predicted peptide MHC binding stability and affinity was associated with antigen immunogenicity. Cancer-testis antigens, endogenous retroviral antigens, and SNV-derived tumor antigens that were immunogenic were found across tumor subclones. By diversifying the immunogenic antigen repertoire beyond SNVs, we achieved nearly 100% tumor subclone coverage, suggesting that broader antigen selection could help immunotherapy target more tumor subclones. In conclusion, this study supports the stability of the immunodominance hierarchy under ICI therapy and a role for broadening antigen discovery to multiple expressional sources in immunotherapy design.

## Introduction

Urothelial carcinoma (UC) is the most common malignancy of the urinary tract, often associated with recurrence following primary therapy and progression to metastatic disease(1–4). In the past, immune checkpoint inhibitor therapy has been used for patients with metastatic and advanced UC who have had disease progression post platinum based chemotherapy or Bacillus Calmette-Guérin (BCG) treatment (5–8). More recently, ICI treatment has been extended into the neoadjuvant, adjuvant, and first-line metastatic settings (9–12). In addition to first-line treatment combinations, ICIs are now being used as maintenance therapy across disease stages in combination with other immunotherapy or chemotherapeutic strategies such as enfortumab vedotin (antibody drug conjugate) with pembrolizumab (anti-PD1) and nivolumab (anti-PD1) plus gemcitibine/cisplatin (6,12,13).

ICIs promote antitumor immunity by blockade of inhibitory signaling pathways that attenuate an anti-tumor T cell response (14). In UC, clinical response to checkpoint blockade has been associated with T cell inflamed tumors, including increased CD8+ T cell infiltration and enrichment of cytotoxic T cell related transcriptional programs (15–20). Patients with tumor infiltrating lymphocytes (TILs) in the tumor microenvironment (TME) have prolonged disease free survival (21–24). However, the breadth of tumor antigen specificities recognized by T cells and how these responses are reshaped by ICI remain poorly understood.

A feature of T cell immunity, including tumor-specific immunity, is T cell immunodominance. An Immunodominance hierarchy is the result of a reproducible ranking of antigen specific T cell responses in order of magnitude to a concentrated set of antigens despite the availability of a broader range of antigenic targets. This phenomenon was first characterized in antiviral immunity, where CD8^+^ T cell receptors were shown to recognize only a small fraction of potential antigens constituting a hierarchy of T cell responses (25–28). Immunodominant antigenic peptides are the subset of predicted or presented peptides that reproducibly elicit the strongest antigen-specific T cell responses within a given system (25,26,28,29). Subdominant antigenic peptides are those that mostly elicit lower frequency T cells (26,30,31). Epitopes within the immunodominant class of the hierarchy are critical to both eliminating virally-infected cells and contributing to the protective memory T cell pool (27,31,32). In cancer, studies using exogenous and engineered antigen systems have demonstrated that T cells specific to the engineered tumor antigen system can also exhibit immunodominance. Checkpoint blockade can alter immunodominance hierarchies, shifting subdominant T cell responses to dominant T cell responses (33–36). In addition, T cell clonal competition across the immunodominance hierarchy may be associated with attenuation of vaccine responses (bioRxiv, 2025.10.26.684631). Because many studies of tumor antigen immunodominance have relied on xenogeneic, engineered, and/or simplified antigen systems, the organization of endogenous tumor antigen-specific responses to physiologically expressed tumor-derived antigenic peptides remains incompletely defined.

In this study, we mapped the tumor antigen immunodominance hierarchy in the BBN963 murine model of basal-like bladder cancer due to its high mutational burden, T cell infiltration, physiologic expression of endogenous tumor-derived antigens, and reflection of human basal like bladder cancer (37). We identified tumor antigens from multiple genomic sources which cover expressional candidate tumor-derived peptides and the diversity of mutational types present (38). We found that the immunodominance hierarchy was stable among individual mice. ICI treatment (anti-PD-1 or anti-CTLA4) did not elicit a considerable shift in immunodominance of CD8^+^ T cell responses to subdominant epitopes, despite previous studies reporting ICI shifting subdominant T cell responses to dominant (35). We analyzed potential determinants of tumor antigenic peptide immunogenicity and found that predicted binding stability and affinity of peptide MHC is associated with immunogenicity. We found the immunodominance hierarchy of antigens discovered in our set of predicted epitopes reached a near 100% coverage of tumor subclones based on single-cell long read sequencing, however SNV derived antigens covered only 65% of tumor subclones indicating broadening antigen class could help target a broader range of tumor subclones.In contrast, SNV-derived antigens were detected in only 65% of tumor-cell subclones. Collectively, these data show diverse endogenous tumor antigen peptides derived from mutations (SNVs, insertion deletions, fusions) but also endogenous cell gene expressed antigens (ERVs and CTAs) elicit an immunodominance hierarchy amongst CD8^+^ T cells that is relatively stable between untreated and ICI treated animals.

## Results

### BBN963 basal-like bladder cancer has a diverse repertoire of tumor antigens

To evaluate if predicted tumor antigen peptides are immunogenic and induce an immunodominance hierarchy of T cells in the BBN963 model of bladder cancer, we predicted antigenic peptides from whole exome and transcriptome sequencing using LENS (38), then we tested predicted antigenic peptides for immunogenicity using a multiplex IFN-γ ELISpot method with post-hoc single peptide validation **(Figure 1)**. One hundred and forty-five predicted tumor antigen peptides were shared in three biological replicates **(Supplemental Figure 1)**, and then assessed for immunogenicity in a matrixed ELISpot design. Results of the matrixed ELISpot were deconvolved, validated in a round of single-antigen ELISpots, and ranked based on IFN-γ production from enriched CD8^+^ T cell populations **(Figure 1).**

**Figure 1.**
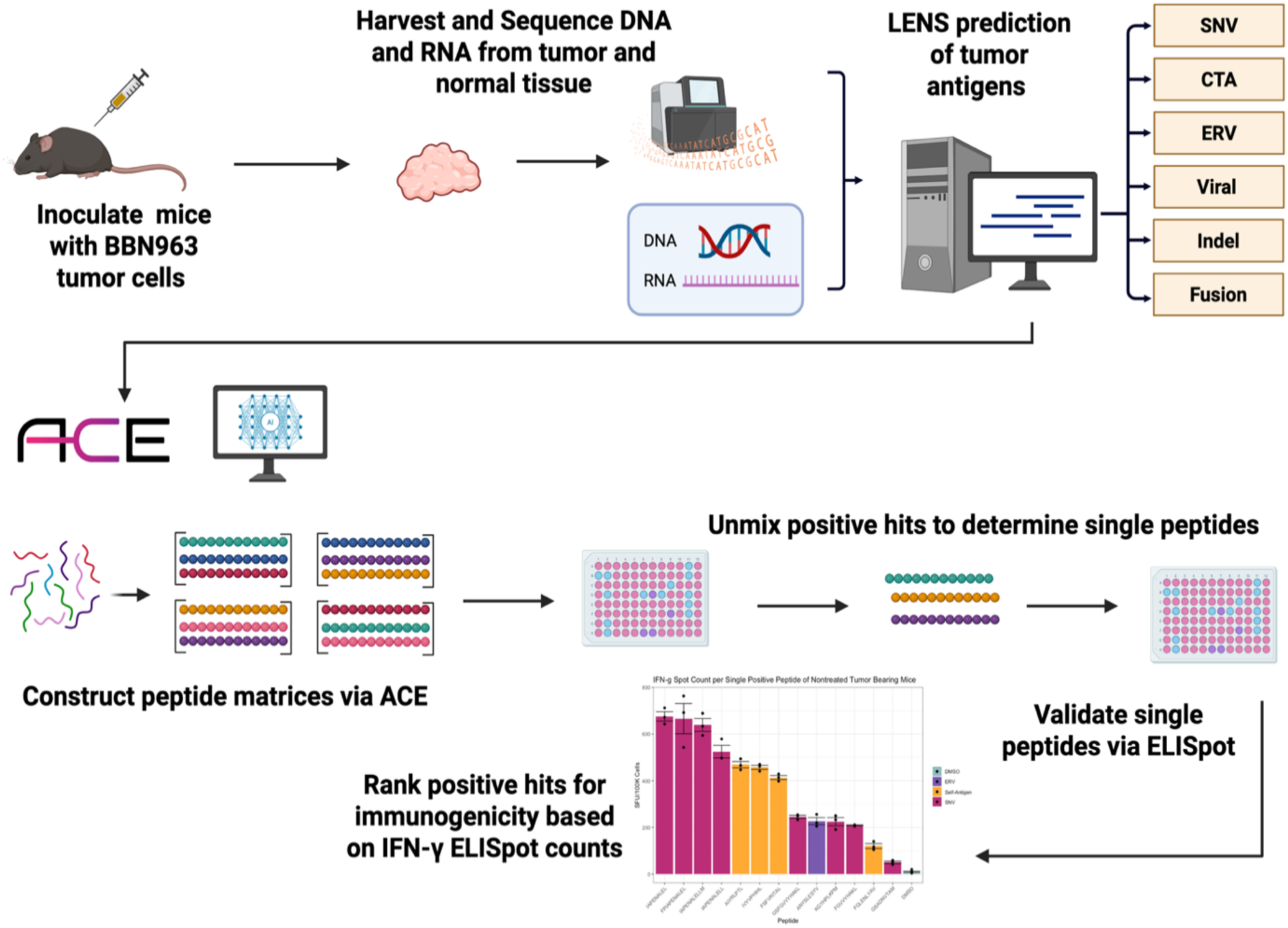
Experimental and computational workflow for discovering the immunodominance hierarchy of CD8^+^ T cells recognizing tumor antigens in the BBN963 murine model of bladder cancer. Three BBN963 tumors were analyzed by whole exome and transcriptome sequencing. Data were analyzed using the Landscape of Effective Tumor antigen Software (LENS) to predict tumor antigens derived from a set of genomic sources (SNVs, indels, fusions, splice variants, viruses, endogenous retroviruses, and cancer/testis antigens). The set of predicted antigenic peptides was evaluated using ACE Configurator for ELISpot (ACE) to construct peptide matrices for large scale ELISpot testing. IFN-y production measured by ELISpot was used to determine relative immunodominance of MHC Class I restricted epitopes recognized by CD8^+^ T cells.

Across the three BBN963 tumors examined, 145 antigenic peptides were predicted to be shared. These mostly comprised SNV, ERV, and CTA-derived peptides **(Figure 2A)**. We next evaluated features of these predicted antigenic peptides by the number of biological replicates in which they were identified. Average RNA read support was higher in tumor antigens found in three replicates versus one or two, while there was no significant difference between antigens found in one or two replicates **(Figure 2B).** Predicted peptide MHC binding stability estimated by NetMHCstabpan1.0 was higher for tumor antigens found in three replicates versus one or two, while there was no significant difference between antigens found in one or two replicates **(Figure 2C)**. Predicted peptide MHC binding affinity was not significantly different between groups **(Figure 2D)**, however of note, 500nM predicted binding affinity was used as a filtering heuristic in LENS such that no tested peptides had predicted binding affinity > 500nM. MHCflurry 2.0 processing score, which is a prediction metric of antigen processing steps that are not MHC-dependent, was significantly higher in antigens found in three replicates versus two or one **(Figure 2E)**, and similarly MHCflurry 2.0 presentation score, which is an allele dependent metric combining binding affinity and processing, was significantly higher in antigens found in three replicates versus two or one **(Figure 2F)** (39). In summary, predicted tumor antigen peptides found in three replicates maintained higher average RNA reads, peptide MHC binding stability, and MHCflurry presentation and processing scores in comparison to tumor antigen peptides found in only one or two replicates. Based on these metrics, we used the core of 145 tumor antigen peptides in our studies of assessing the T cell immunodominance hierarchy to BBN963 bladder cancer.

**Figure 2.**
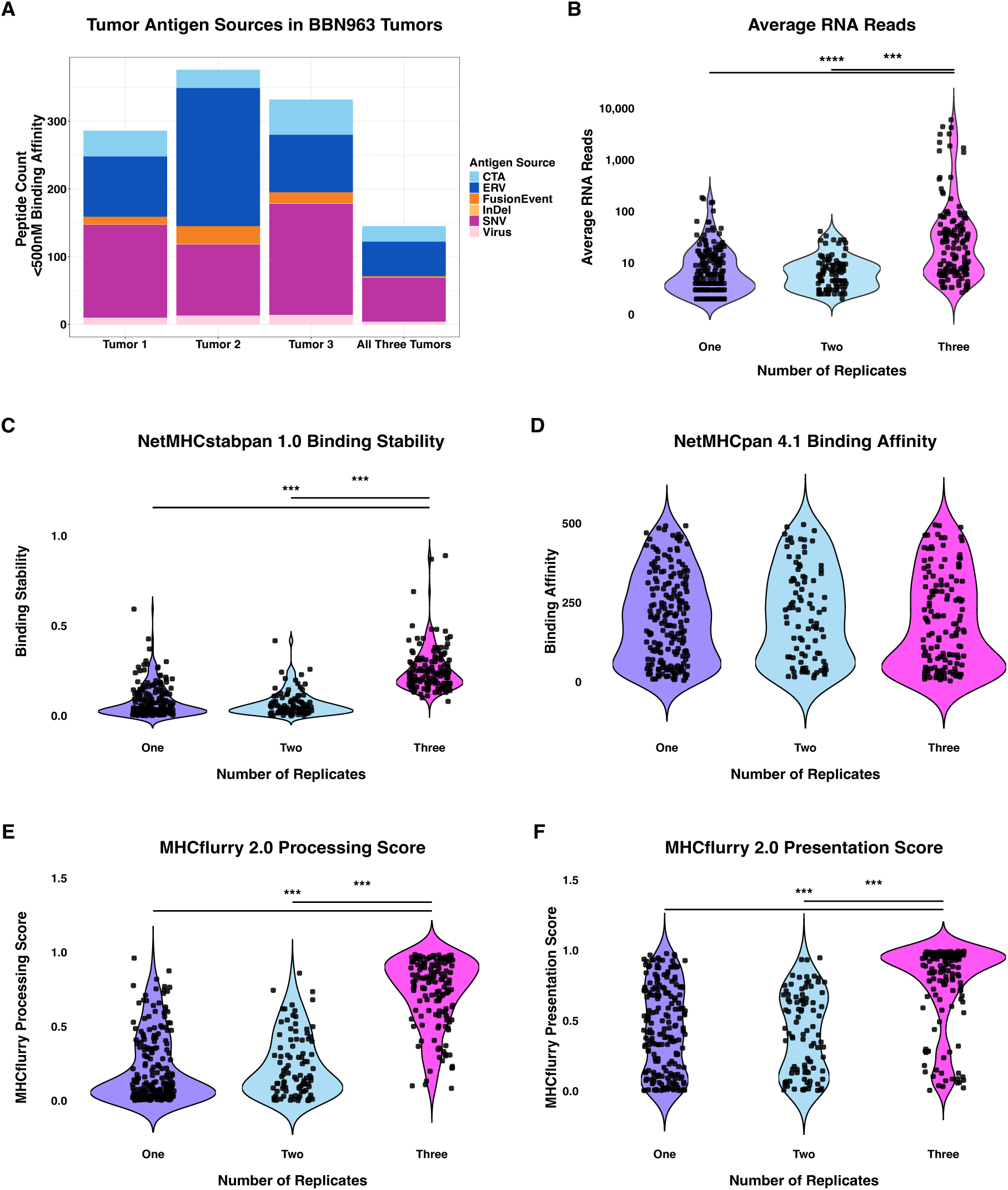
Features of predicted antigenic peptides in BBN963 amongst biological replicates. (A) Predicted tumor antigen peptide count, genomic source per replicate, and number of antigen peptides shared amongst three biological replicates (B) RNA-seq read support for antigen peptide coding transcripts (C) Predicted binding stability (netMHCStabPan) (D) Predicted binding affinity (NetMHCpan4.1) (E) MHCFlurry processing score (F) MHCFlurry presentation score of all predicted antigenic peptides found in one, two, or three replicates. One-way Anova with Tukey’s HSD test conducted. (P < 0.001 ***, P < 0.01 **, P < 0.05 *).

### Immunogenic tumor antigens form a stable immunodominance hierarchy

We conducted high-throughput matrixed ELISpots via peptide pooling in untreated tumor-bearing animals and saw a distribution of ELISpot responses that indicated preferential T cell responses to peptide pools **(Figure 3A)**. When validated immunogenic peptides were mapped back to peptide pools, the distribution of positive peptide pools had higher T cell IFN-γ responses in comparison to negative pools **(Figure 3A)**. Deconvolution of matrixed ELISpot via ACE identified 19 peptides spanning SNV, ERV and CTA antigens that were next tested individually. We classified antigens that elicited strong and frequent CD8⁺ T cell responses with statistical significance measured by distribution free resampling with 2x above negative control (DFR 2x) as immunogenic **(Supplemental File 1)** (40,41). This yielded 18 of our 19 validated antigen peptides as immunogenic. Immunogenic responses varied by mutational class and predicted MHC restriction and included non-mutated antigen classes, such as CTAs and ERVs **(Figure 3B)**. Notably, the strongest T cell responses were directed against an SNV-derived tumor antigen (IAPENALEL) predicted to bind H2-Db. T cell responses were also found to its overlapping peptide variants (IAPENALELL, IAPENALELLM, FPIAPENALEL). In addition, another core antigen and its variant (FGVVYHAKL and GSFGVVYHAKL) were predicted to bind to H2-Kb and elicited subdominant T cell responses **(Figure 3C).** We calculated peptide biochemical similarity of immunogenic antigenic peptides to determine if immunodominant and subdominant antigens shared similar biochemical properties. Peptides sharing core sequences and thus biochemical properties clustered tightly together as shown in the cladogram (42) and shared immunodominance tier **(Figure 3C).**

**Figure 3.**
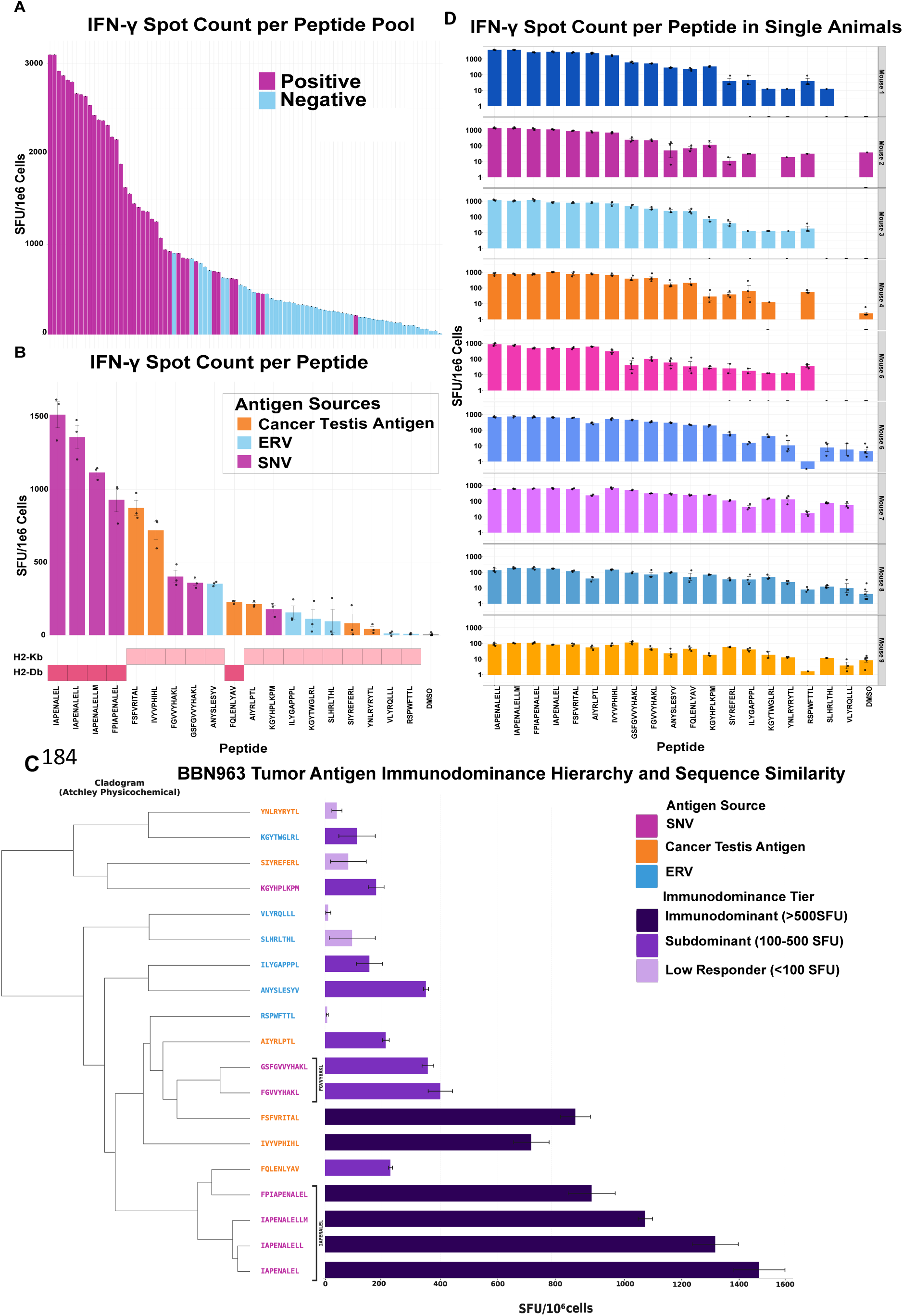
The immunodominance hierarchy in BBN963 is stable amongst mice. 6-week-old C57BL/6 female mice were inoculated with 3 x 10^6^ BBN963 cells, sacrificed at 10 days post tumor inoculation, and assessed for tumor antigen-specific CD8^+^ T cell responses via IFN-y ELISpot in pooled animal studies n=15 and individual animal studies (biological replicates n=9). (A) IFN-y spot count of matrixed peptide pools in pooled animal studies. Peptide pools containing positive antigens are shown in magenta and pools that are negative shown in light blue. (B) Single peptide validation of tumor antigens via ELISpot. Tumor antigens are mapped to genomic source and predicted MHC restriction (H2-Kb and H2-Db). (C) Immunodominance hierarchy of Tumor Antigens across immunogenic peptides with peptide biophysical similarity cladogram displaying ELISpot response to peptides with same core sequences. (D) Validation of immunodominance hierarchy of immunogenic peptides in single animal studies.

When further exploring what biochemical factors may contribute to immunodominant and subdominant groups, we found secondary structure was significantly different across groups (KW p = 0.039; Dunn-Holm p = 0.035), and immunodominant peptides were significantly enriched in amino acids with high helical structure favorability (Spearman ρ = −0.554, p = 0.014; Kruskal-Wallis p = 0.039; Dunn-holm p = 0.035) **(Supplemental Figure 2)**. Other biochemical factors such as polarity/hydrophobicity, molecular size, codon composition, and electrostatic charge were not significantly different across the immunodominance tiers.

To see if the hierarchy of immunogenic tumor antigen peptides was stable across individual animals, single animal ELISpots were conducted to validate predicted tumor antigens **(Figure 3D)**. A total of nine animals from two independent experiments, we observed a relatively stable hierarchy of immunonogenic peptides.

### The BBN963 immunodominance hierarchy remains stable under anti-PD-1 and anti-CTLA4 treatment

To determine if ICI would shift T cell immunodominance hierarchies, we repeated our matrixed IFN-γ ELISpot experiments on animals treated with anti-PD-1 or anti-CTLA4 inhibitory monoclonal antibodies. ICI treatment was given biweekly for two weeks, after which T cell responses were assessed via matrixed ELISpot seven days after the last treatment **(Figure 4A)**. In comparison to mice treated with PBS and isotype control antibodies, anti-PD-1-treated animals had increased T cell responses to peptide pools **(Figure 4B).** Each treatment arm was deconvoluted and single peptide validation was conducted via IFN-γ ELISpot. Overall, anti-PD-1-treated animals had higher IFN-γ responses to tumor antigens in comparison to control arms while the relative hierarchy of antigens was maintained to immunogenic peptides **(Figure 4C)**. Similarly, anti-CTLA4-treated animals had increased T cell responses to peptides pools in comparison to controls **(Figure 4D).** As with anti-PD-1 treatment, the IAPENALEL peptide and its variants remained immunodominant and the relative hierarchy remained stable in comparison to untreated animals **(Figure 4E)**.

**Figure 4.**
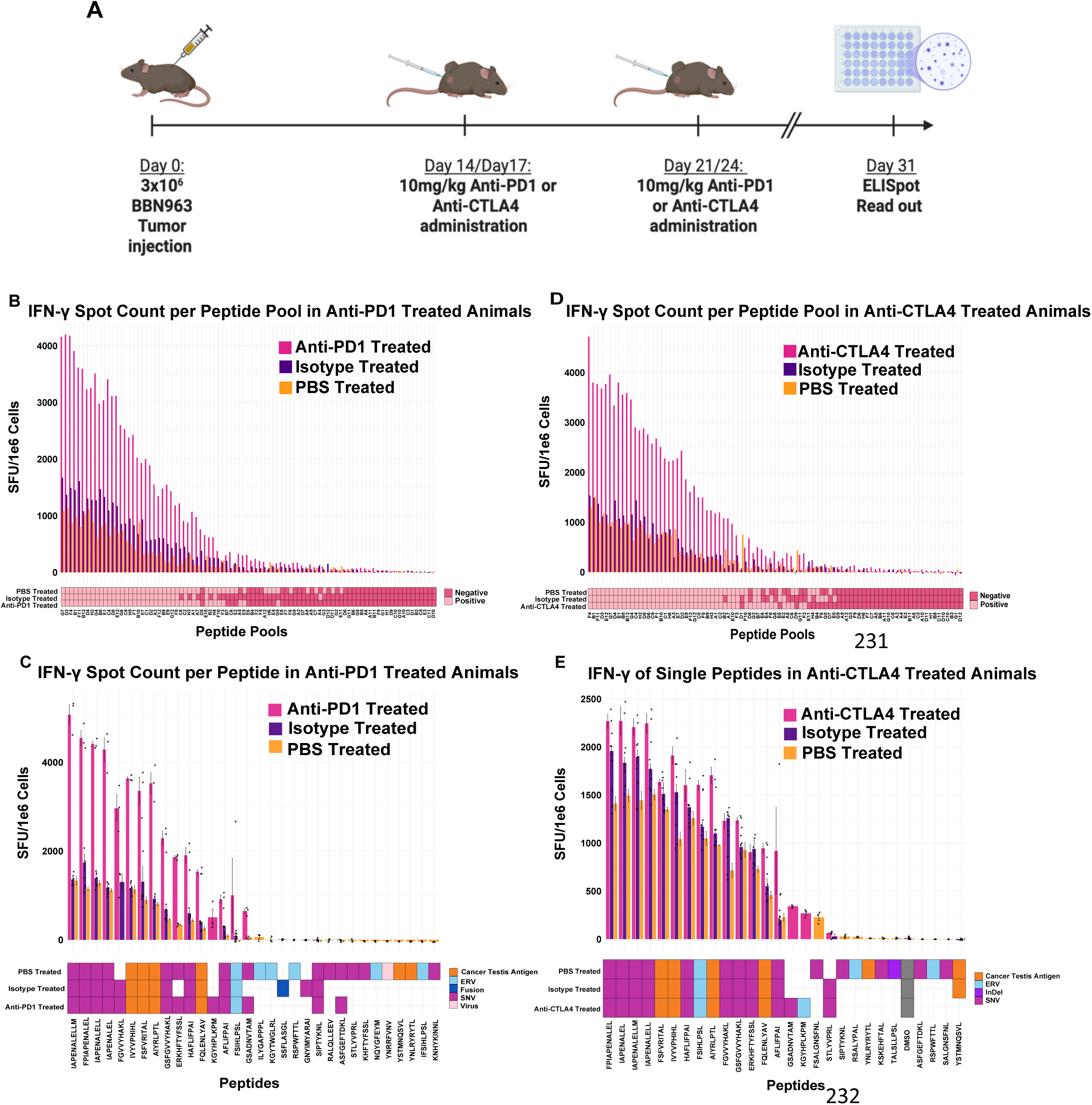
Immunodominance hierarchy is relatively conserved under anti-PD1 and anti-CTLA4 monotherapy. (A) Experimental design: 6-week-old C57BL/6 female mice were inoculated with 3 x 10^6^ BBN963 cells. Assessment of antigen-specific CD8^+^ T cell responses was done as outlined in figure 1 (n=10 per treatment arm). (B) 145 peptide matrixed IFN-y ELISpot results of anti-PD1 treated tumor-bearing animals as deconvoluted by ACE. (C) Single peptide validation of anti-PD1 treated animals showing the genomic source of each antigen. (D) 145 peptide matrixed IFN-y ELISpot results of anti-CTLA4 treated tumor bearing animals as deconvoluted by ACE. (E) Single peptide validation of anti-CTLA4 treated animals showing the genomic source of each antigen.

### Immunopeptidomics validation of immunodominant tumor antigens

To test MHC presentation of predicted antigenic peptides and to complement our immunogenicity experiments, we performed shotgun mass spectrometry (MS) immunopeptidomics studies on the tumor cell line via two independent labs designated as Protocol 1 (Jaeger Lab) and Protocol 2 (Biognosys). From our set of 145 predicted antigenic peptides, protocol 1 detected fourteen tumor antigens and protocol 2 detected four **(Figure 5A)**. Three of the tumor antigens detected by protocol 1 were immunodominant, including the SNV-derived IAPENALELL and its variant IAPENALELLM, along with the CTA-derived peptide AIYPRLPT **(Figure 5A)**. The MS/MS spectra of both IAPENALELL **(Figure 5B)** and AIYPRLPT **(Figure 5C)** from protocol 1 showed consistent b- and y- ion fragmentation for the two antigens, supporting their identification by MS. Of the 145 predicted antigenic peptides tested for immunogenicity in untreated animal studies, 3 were found to be immunogenic and detected by MS (2% of peptides tested) while 11 were detected by MS but not found to be immunogenic (8% of peptides tested) **(Figure 5D)**. In addition, 15 of the predicted tumor antigens were immunogenic but not detected by MS constituting 10% of the 145 tumor antigen peptides tested **(Figure 5D)**. When compiling immunogenicity across untreated and ICI-treated experimental groups, the number of MS-detected and immunogenic peptides remained the same, while the number of immunogenic antigenic peptides not detected by MS increased and constituted 14% of the peptides tested **(Figure 5E).**

**Figure 5.**
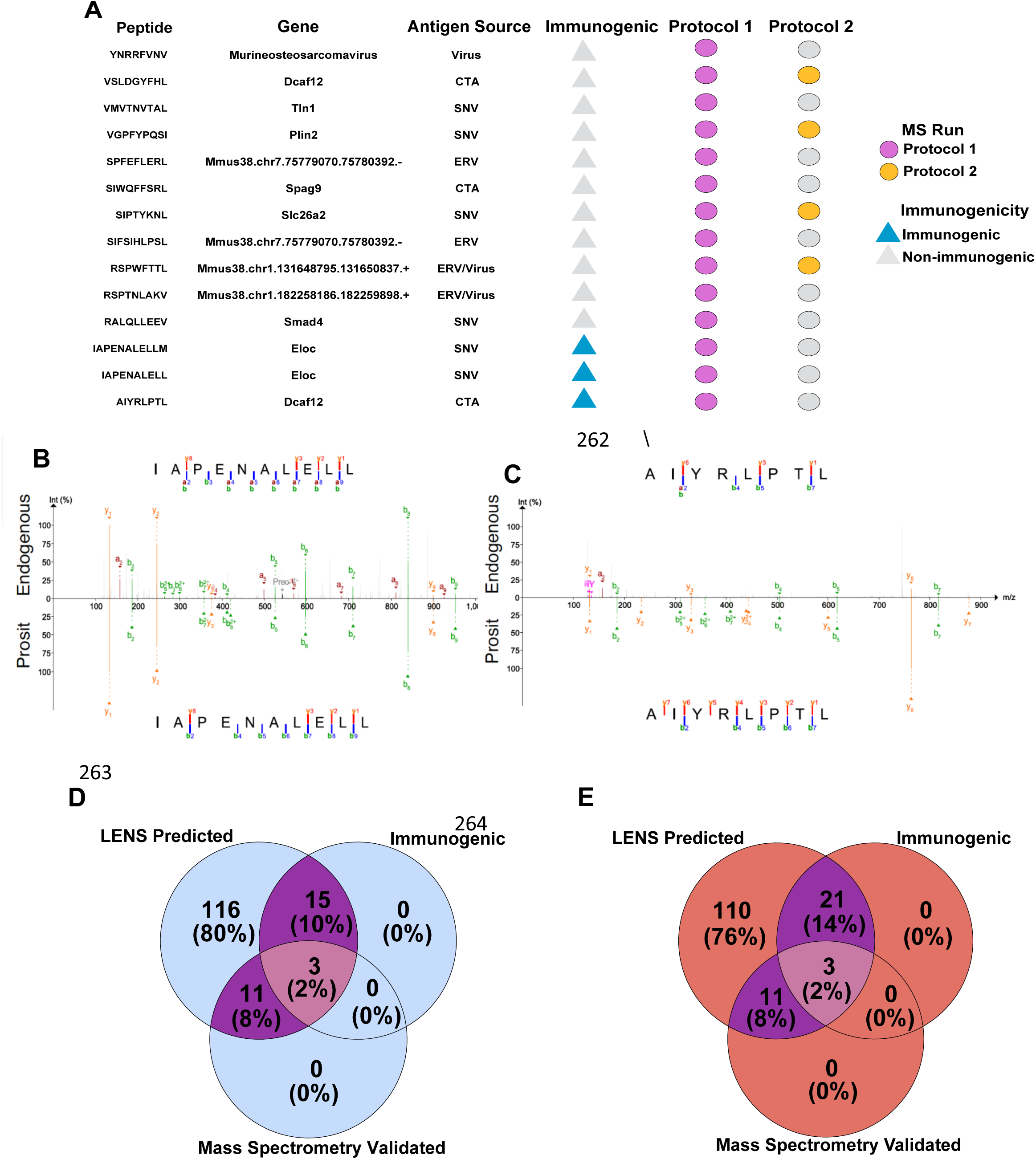
Immunopeptidomics validation of predicted BBN963 predicted peptides, (A) Predicted tumor peptides discovered by mass spectrometry immunopeptidomics, showing their coding gene, genomic source type, and immunogenicity status based on matrixed ELISpot studies. (B) Mirror plot showing the endoengous MS/MS spectrum of immunogenic antigen IAPENALELL compared to the Prosit predicted spectrum. (C) Mirror plot showing the endoengous MS/MS spectrum of immunogenic antigen AIYRLPTL compared to the Prosit predicted spectrum. (D) Venn diagram of the 145 shared BBN963 predicted tumor peptides found to be immunogenic in untreated tumor-bearing animals and identified by immunopeptidomics. (E) Venn diagram of the 145 shared BBN963 predicted tumor peptides found to be immunogenic in untreated and ICI-treated animals and identified by immunopeptidomics. N=4 technical replicates in Protocol 1, N=3 technical replicates in Protocol 2. Antigens were considered immunogenic based on DFR 2x statistical testing with p < 0.05.

### Peptide MHC binding stability and affinity associates with immunogenicity

To better understand tumor antigen features associated with immunogenicity, we performed regression analysis considering predicted peptide MHC binding stability, predicted peptide MHC binding affinity, MHCFlurry processing and presentation scores, average RNA read support, and detection by MS as predictor variables and immunogenicity as the response variable. Immunogenicity was encoded as a binary variable and considered a rare event in our dataset. In univariable logistic regression, peptide MHC binding stability (OR = 51.22, 95% CI: 1.862-1409.3, p = 0.02) and peptide MHC binding affinity (OR = 0.382, 95% CI: 0.153-0.951, p = 0.039) were associated with higher odds of immunogenicity **(Figure 6A and Supplemental Table 1).** To determine whether predictor variables had an association with immunogenicity while adjusting for the effects of other covariates, we used a multivariable firth model as a penalized logistic regression to handle rare positive events and mitigate small sample bias. In this model, predicted peptide MHC binding stability (OR = 1.556, 95% CI: 1.019-2.375, p = 0.041) was the only variable associated with higher odds of immunogenicity in the presence of other covariates **(Figure 6B and Supplemental Table 2).** When assessing firth model performance, we found the area under receiver operating curve (ROC) measure to be modest (Apparent ROC-AUC = 0.703; LOOCV ROC-AUC = 0.558), but when cross validated it is barely is above 0.5 for random ranking **(Figure 6C)**. Because immunogenic peptides were rare, 18 events total out of 145, we evaluated the precision-recall curve AUC (PR-AUC) for model performance. Apparent AUC of PR was 0.36 and the LOOCV AUC was 0.209, modestly above baseline of 0.125 indicating limited recall for immunogenic peptides **(Figure 6C)**.

**Figure 6.**
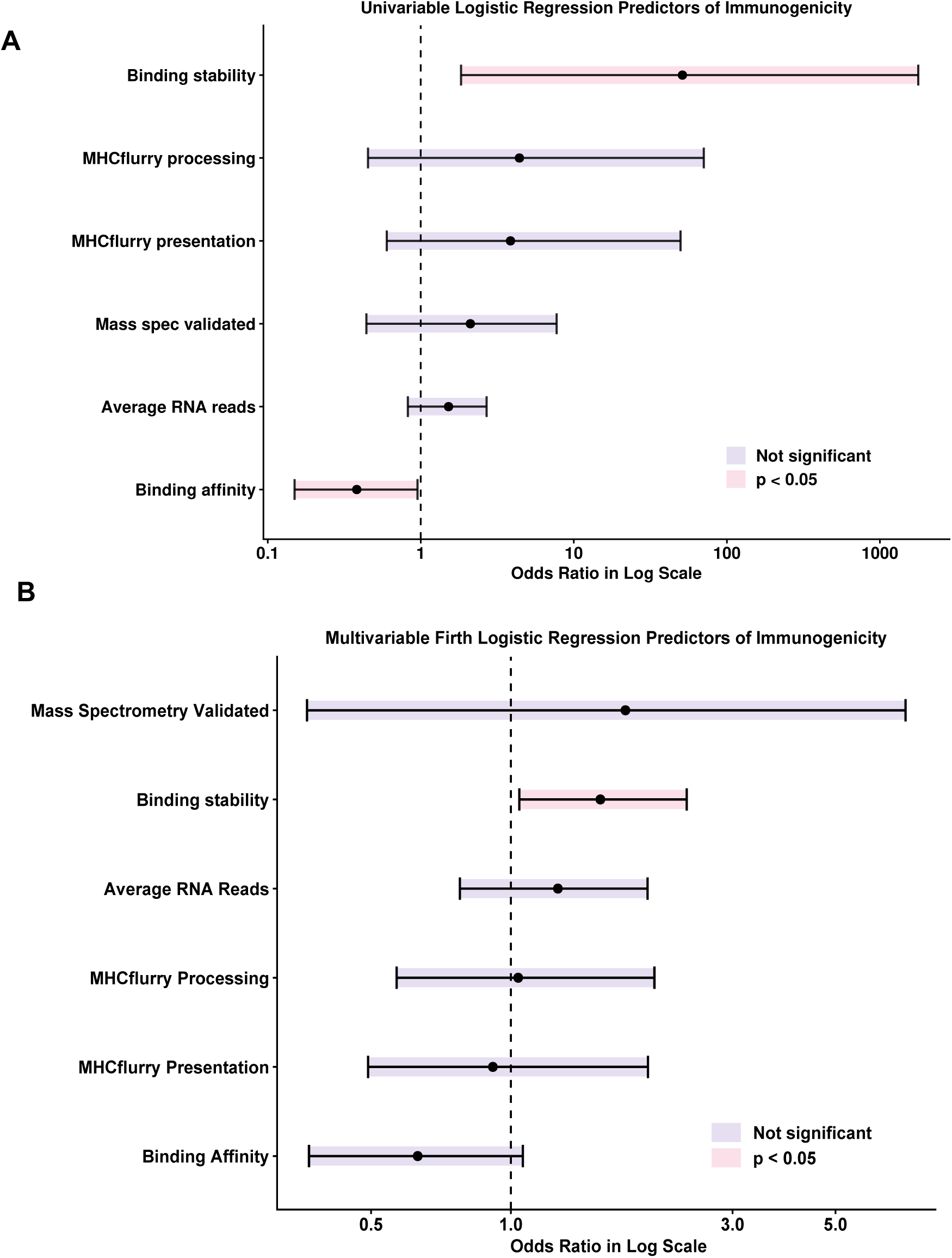
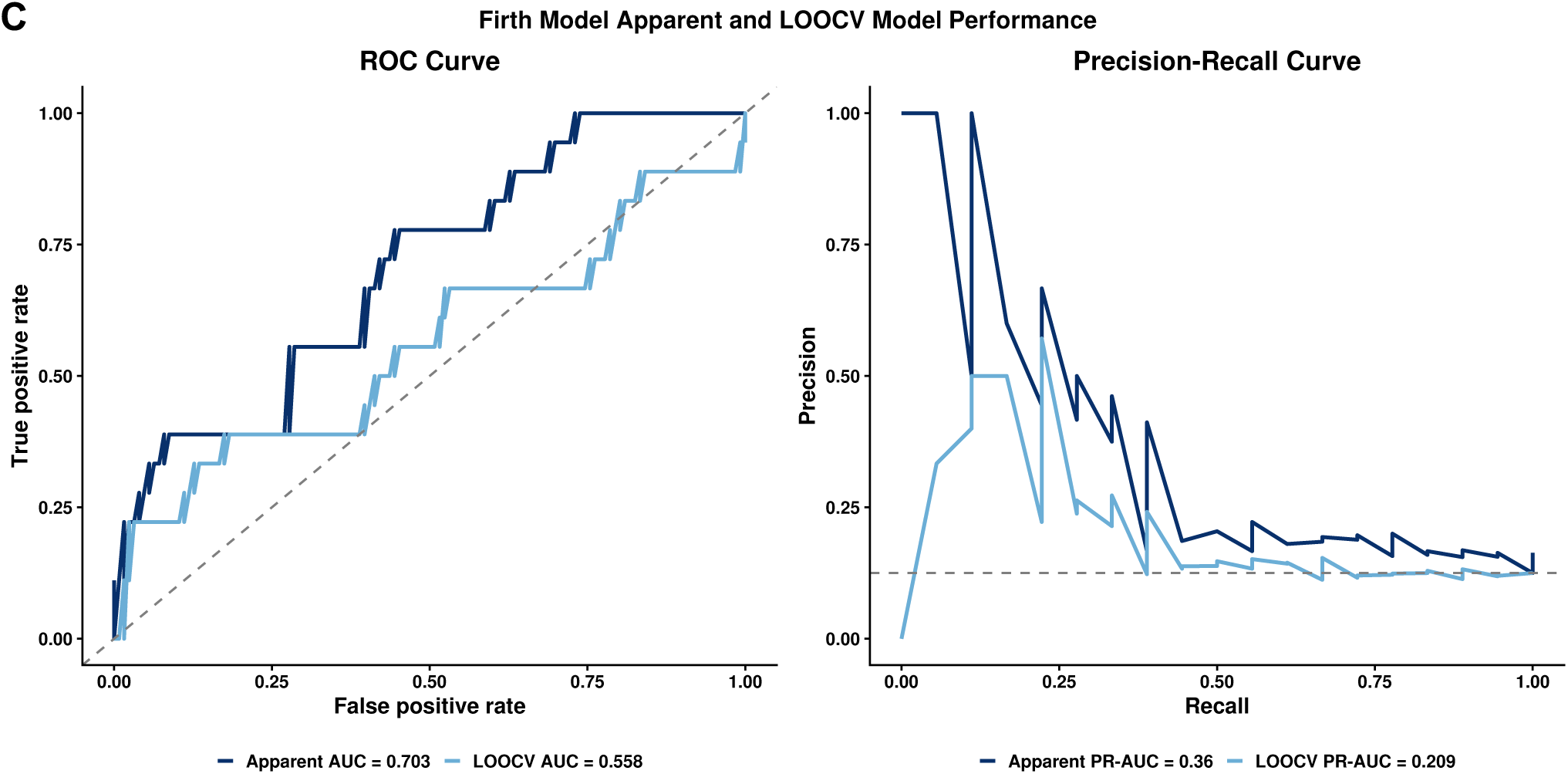
Univariable and Multivariable Modeling of Immunogenicity in the BBN963 Model of Bladder Cancer. (A) Odds ratio of tumor antigen feature associations with immunogenicity via univariable logistic regression. Dotted line is 1 at null value, representing no effect on immunogenicity. Lines represent 95% confidence interval (CI), and colored bars indicate if features were statistically significant (p-value < 0.05). (B) Odds ratio of predicted antigenic peptide metrics in Firth multivariable model. Dotted line is 1 at null value, representing no effect on immunogenicity. Lines represent 95% confidence interval (CI), and colored bars indicate if features were statistically significant (p-value < 0.05). (C) Firth Model Performance curves. ROC and precision recall (PR) curves with apparent vs leave-one-out cross-validation (LOOCV) AUC.

### Validated immunogenic antigenic peptides cover nearly all BBN963 tumor-cell subclones

To determine the heterogeneity of tumor antigen peptide expression among tumor-cell subclones, we developed a tumor antigen profile network data structure generated from long read single cell transcriptomics data **(Figure 7A)**. Each node in the network represents a tumor antigen peptide profile, and profiles that differ by a single tumor antigen peptide are linked by an edge. **Figure 7A** details a tumor antigen network showing heterogeneity of expressed tumor antigen peptide profiles from our list of peptides that are immunogenic and mass spec validated. The proportion of tumor cells expressing validated immunogenic tumor peptides nearly reaches 100% **(Figure 7B)**. Across SNV-derived tumor antigen peptides, IAPENALEL and its variants covered nearly 40% of all BBN963 cells while cumulatively the immunogenic SNVs cover over 60% of the BBN963 cells **(Figure 7C)**. Across CTAs, we saw a cumulative coverage of around 80% while tumor antigen peptides FQLENLYAV and IVYPHIHL had the highest coverage **(Figure 7D)**. Across ERV tumor antigen peptides, a cumulative coverage across all antigens was 85%, driven by tumor antigen peptide ILYGAPPPL which was found to have the highest coverage above 80% **(Figure 7E)**. From our immunogenic and MS validated tumor antigen peptides, we found cumulative coverage across BBN963 cells to be about 45% **(Figure 7F)**.

**Figure 7.**
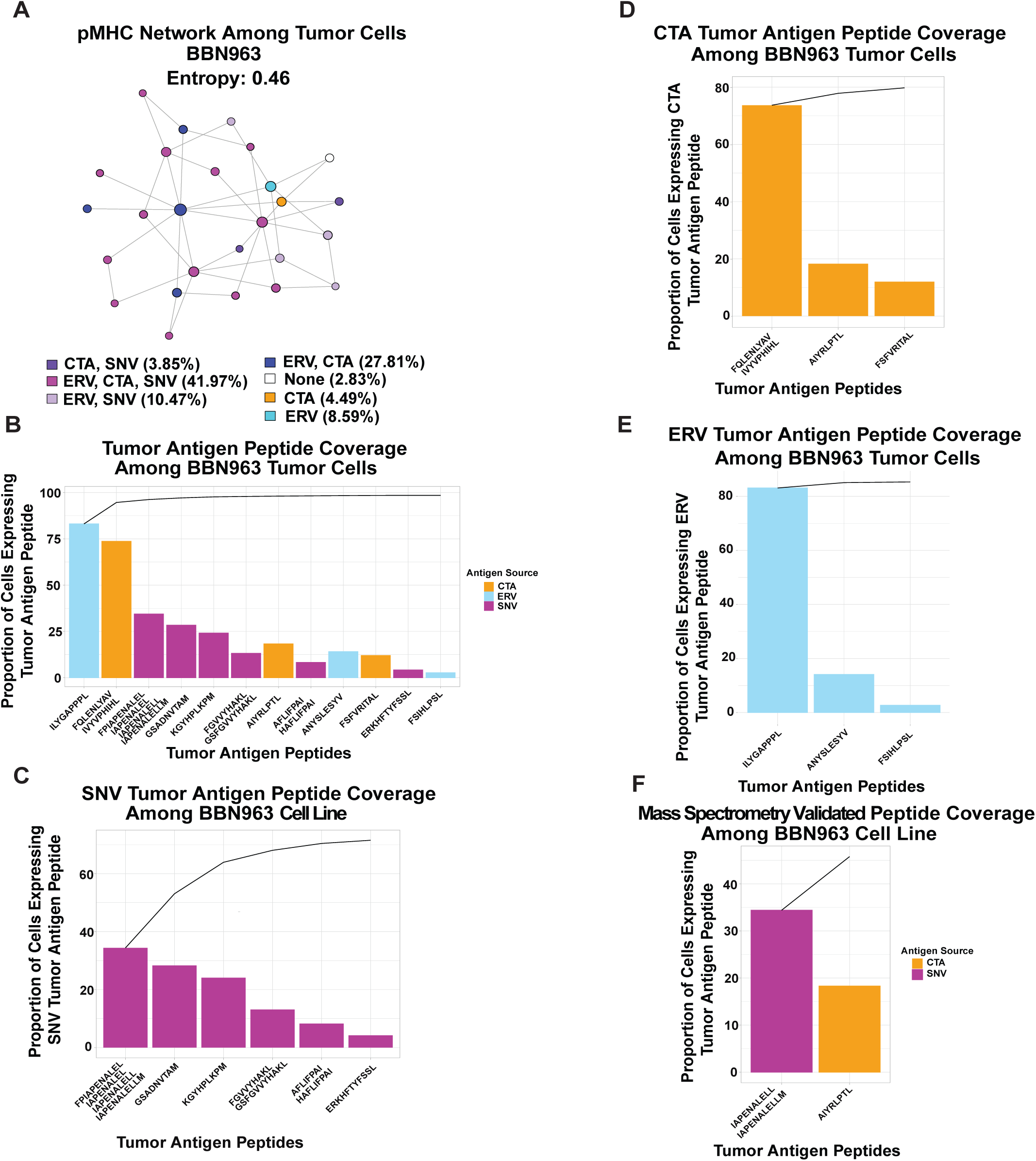
Tumor antigen network profiling expression and coverage of tumor antigens across BBN963 single cells. (A) Tumor antigen network depicting proportion of cells sharing tumor antigen peptide expression profile based on immunogenic and mass spec validated tumor antigens. (B) Proportion of cells expressing immunogenic peptides. Line depicts coverage of each antigenic peptide across tumor cells. (C) Proportion of cells expressing immunogenic SNV’s. Line depicts coverage of antigenic peptides across tumor cells. (D) Proportion of cells expressing immunogenic CTA’s. Line depicts coverage of each antigenic peptide across tumor cells. (E) Proportion of cells expressing immunogenic ERV’s. Line depicts cumulative coverage of antigenic peptides across tumor cells. (F) Proportion of cells expressing immunogenic and mass spec validated immunogenic peptides. Line depicts coverage of antigenic peptide across tumor cells.

## Discussion

A central finding of this study is that from 145 BBN963 predicted tumor antigen peptides tested, only 18 of the 19 validated antigen peptides were immunogenic based on stastistical significance as determined by DFR 2x (40,41) **(Supplemental File 1).** These generated a stable immunodominance hierarchy across untreated tumor bearing animals. Despite each animal being able to theoretically generate a highly diverse T cell receptor repertoire through V(D)J recombination, antigen-specific T cell responses were consistent in our model. This bias has been reflected in the public TCR space in the viral, autoimmune, and tumor-associated antigen contexts where individuals share the same or similar TCR’s to the same antigenic epitope (43–48). This phenomenon may reflect biases in TCR recombination, thymic selection, or precursor frequency as well structural constraints in TCR and peptide-MHC interactions. Thus, the stability of the BBN963 immunodominance hierarchy across animals suggests that TCR recognition of certain peptide-MHC complexes repeatedly recruits clonotypes that are biased toward generation by V(D)J recombination, thymic positive selection, and expansion post-antigen stimulation.

In line with this, the most immunodominant antigen was an SNV-derived peptide (IAPENALEL), rather than antigen peptides with more distinct from self amino acid sequences such as ERV, InDel, or fusion classes. This suggests that immunodominance is not solely driven by antigen class but likely emerges from a combination of T cell intrinsic and extrinsic factors such as TCR clonotype, TCR affinity and stability to peptide-MHC, T cell state, antigen abundance, MHC binding stability, antigen processing, and precursor T cell availability. Prior work in antigen-engineered tumor models, vaccine design, and the viral immunodominance space has shown that immunodominance can be established with multiple competing T cell responses, and antigens with stronger binding stability to MHC drive dominant responses (34,49–51)(bioRxiv, 2025.10.26.684631). Future studies using peptide/MHC multimer sorting followed by TCR sequencing would help determine the extent to which antigen-specific clonotypes are repeatedly generated across animals, whether dominant and subdominant responses differ in TCR diversity, and whether dominance reflects precursor frequency, clonal expansion, or functional T cell state.

We also found that the immunodominance hierarchy was maintained under ICI treatment. We initially hypothesized that anti-PD-1 or anti-CTLA-4 therapy might shift the immunodominance hierarchy by enhancing subdominant antigen-specific T cell populations to become dominant. Instead, the relative hierarchy of CD8^+^ T cell responses was preserved. Checkpoint blockade acts through distinct cellular mechanisms that depend on T cell state and preferentially expands T cells that are more effector/exhausted-like (52). Our findings suggest that our immunodominant and subdominant T cell populations may retain a more effector/exhausted-like state, making them more sensitive to ICI treatment. However, this needs to be verified with follow up flow cytometry studies using peptide/MHC multimers to analyze the phenotype of antigen-specific T cells. In addition, these results differ from prior work showing that neoadjuvant anti-PD-1 therapy can reverse immunodominance in a mouse oral carcinoma model containing the defined OVA xenoantigen and p15E retroviral antigen (35). Together, these studies suggest that ICI-mediated remodeling of immunodominance may be highly context-dependent.

Most predicted antigenic peptides in our dataset were not shown to be immunogenic by IFN-γ ELISpot. While computational tumor antigen prediction generates a plausible list of tumor antigens, the majority of those antigens are either not naturally presented, not recognized by T cells, or not capable of inducing a T cell response. Benchmarking studies have shown that a minority of predicted epitopes are experimentally immunogenic and improved prioritization requires integration of features related to antigen presentation and T cell recognition (49,53,54). To further validate our tumor antigens, we conducted mass spectrometry-based immunopeptidomics. In our study, 14 of the 145 predicted antigenic peptides were MS-validated, and three of these MS-validated peptides generated reproducible CD8^+^ T cell responses. We then proceeded to model determinants of immunogenicity in our dataset to see what antigen intrinsic features were associated with T cell response. We conducted univariable and multivariable modeling of peptide features to test for associations with immunogenicity. We found predicted peptide MHC binding stability and peptide MHC binding affinity were associated with immunogenicity, whereas RNA reads, MS validation, and predicted processing or presentation scores were not significant predictors in this dataset. Because our dataset contained few immunogenic antigens, we interpret these results as exploratory and hypothesis-generating.

We also observed that multiple classes of tumor antigen peptides were needed to approach broad coverage of BBN963 tumor cell tumor-cell subcloness. SNV-derived immunogenic peptides alone covered approximately 60–65% of BBN963 cells. Inclusion of additional antigen sources, including CTA- and ERV-derived antigen from our validated immunogenic peptides, increased coverage to nearly 100% across tumor-cell subclones. Tumor heterogeneity can limit the effectiveness of therapies directed against a small number of antigens, particularly when those antigens are unevenly distributed across tumor cell populations (55). Our results demonstrate that immunotherapeutic strategies could benefit from incorporating broader classes of tumor antigens beyond SNV’s to capture larger fractions of tumor heterogeneity and increase therapeutic efficacy.

Our study has several limitations. First, this study was performed in a single murine cancer model. While BBN963 is a useful model for studying bladder cancer with a high mutation burden and T cell infiltration, the antigen landscape, T cell repertoire, and response to ICI may differ in other tumor models (37). Second, our immune monitoring relied primarily on ELISpot analysis. ELISpot is able to detect IFN-γ-producing antigen-specific T cells, but it does not capture all aspects of T cell function, including phenotypic state, cytotoxic capacity, proliferation, other cytokine production. Third, we did not map T cell population-level antigen specificity to TCR sequence and T cell phenotype. As a result, the mechanisms underlying hierarchy stability, such as clonotype diversity, competition, and state, remain unresolved. Finally, because the number of immunogenic peptides was small, statistical modeling results should be validated in larger datasets.

In summary, our study demonstrates that an immunodominance hierarchy comprised of endogenous tumor antigens is stable across animals and ICI therapy. We used a scalable high throughput ELISpot assay to assess immunogenicity and determined that ICI therapy can preserve the immunodominance hierarchy. If shifting the immunodominance hierarchy is desired for therapeutic benefit, other approaches such as tumor antigen vaccination or TCR-engineered T cell therapy should be considered. For many tumors, antigens selected from the set of small somatic mutations will fail to cover all tumor tumor-cell subclones; in this case, therapies restricted to that set could drive immune escape and failure to achieve clinical responses. Similarly, therapies targeted at antigens that are predicted but not actually presented by MHC on the tumor cell surface will fail to deliver positive clinical results. Going forward, improvements in sensitivity and accessibility of mass spectrometry-based validation and high-throughput immune monitoring assays will allow us to more deeply explore how immunotherapy shapes the immunodominance hierarchy and its therapeutic implications.

## Materials and Methods

### Mouse Sourcing and Housing

6-8 week old female C57BL/6J wild type mice were ordered from Charles River (Charles River, Cat No 027). All experimental arms were gender and age matched. All studies were performed under an animal protocol approved by the University of North Carolina at Chapel Hill Institutional Animal Care and Use Committee under the Division of Comparative Medicine. Mice were assessed for morbidity according to animal protocol and humanely sacrificed under CO_2_ euthanasia and a secondary method.

### Cell Line Culturing

BBN963 bladder cancer cells were generated as previously reported (37). BBN963 cells were cultured in Gibco DMEM (4.5 g/L D-glucose, L-glutamine, Phenol Red; Cat No 11995-065) supplemented with 10% fetal bovine serum (FBS) and 1x penicillin/streptomycin. DC2.4 cells were cultured in RPMI-1640 (Gibco, Cat No 11875-093) supplemented with 10% FBS, 1x penicillin/streptomycin, 2 mM L-glutamine, 1x nonessential amino acids, 1 mM sodium pyruvate, 55 uM beta-mercaptoethanol, and 1 M HEPES and 0.2 um sterile filtered. All cell lines were maintained at 37 °C in 5% CO_2_.

### Murine BBN963 Tumor Injection

BBN963 were grown to 50 to 70% confluency on day of injection. Cells were trypsinized using 0.05% trypsin-EDTA (Gibco, Cat No 25300-054), washed and resuspended in 1x DPBS (Gibco, Cat No 14190-144) at 3 x 10^6^ cell per 50 uL, then supplemented with 50 uL of Matrigel (Corning, Cat No 354248) to achieve 100 uL injection volume containing 1:1 tumor cells to Matrigel. BBN963 cells were then injected subcutaneously in the flank in 6 to 8 week old C57BL/6J mice under isoflurane anesthesia.

### BBN963 Tumor Induction and Prep for WES and Bulk-RNA-seq

Three BBN963 tumors in 6 week old C57BL6/J mice were harvested 5 weeks post-induction and dissociated using gentleMACS C tubes (Miltenyi Biotec, Cat No 130-093-237) on protocol spleen dissociation program 1 two times in 5-10 mL of complete RPMI-1640 containing 10% FBS and 1x penicillin/streptomycin. Samples were processed through a 70-um filter. Cells were pelleted at 1200 rpm for 5 min at 4 °C, resuspended in 1-5 mL of ACK Buffer (Gibco, Cat No A10492-01), and incubated for 2 min. 20 mL of complete RPMI-1640 were added to cells and pelleted as stated above. Cells were washed in 1x DPBS then placed into Qiagen Allprep DNA/RNA (Cat No 80204) RLT buffer with 2-mercaptoethanol.

### Stranded mRNA-seq Library Preparation and Sequencing

100ng of total RNA was converted into strand-specific mRNA-seq libraries using Illumina’s stranded mRNA prep ligation kit according to manufacturer’s instructions (Illumina, Cat No 20040553). Briefly, oligo(dT) magnetic beads were used to capture mRNAs with polyA tails followed by clean up, fragmentation and denaturation of mRNA. Synthesis of first strand cDNA was conducted with second strand cDNA synthesis performed to generate blunt-ended dsDNA fragments. dUTP chemistry was performed to quench second strand amplification achieving strand specificity. 3’ adenylation was performed on blunt fragments for dual indexed adapter ligation. Libraries were PCR amplified and enriched using magnetic bead-based clean up. Final libraries were quantified and size checked via Qubit 4 Fluorometer using the Qubit dsDNA kit (Thermofisher, Cat No Q32853) and TapeStation 2200 with High Sensitivity D1000 reagents (Agilent, Cat No 5067-5585, and ScreenTape, Cat No 5067-5584) per manufacturer instructions. The size distribution of fragment libraries was between 300-400 bp. Paired-end sequencing was performed using NovaSeq 6000SP (2 x 100 paired end chemistry).

### Whole-exome sequencing (WES) library preparation

Genomic DNA libraries for WES were prepped with Illumina DNA Prep with Enrichment Kit (Cat No 2002553). Briefly, 100 ng of gDNA per tumor was tagmented and ligated with adapter sequences. Post-tagmentation barcoded gDNA was generated via PCR amplification with indexed adaptors. Libraries were cleaned up using a bead-based method with magnetic column separation (AMPure XP Beads). Sample concentration and tagementation success were checked using Qubit 4 Fluorometer Qubit dsDNA kit (Thermofisher, Cat No Q32853) and TapeStation 2200 with High Sensitivity D1000 reagents (Agilent, Cat No 5067-5585 and ScreenTape Cat No 5067-5584) per manufacturer instructions. Libraries were then pooled and probes were hybridized and captured for enrichment of coding exome sequences. Enriched libraries were PCR amplified and cleaned up using AMPure XP Beads and again checked for concentration and fragment size using a Qubit 4 Fluorometer (Qubit dsDNA kit, Thermofisher, Cat No Q32853) and TapeStation 2200 with High Sensitivity D1000 reagents (Agilent, Cat No 5067-5585 and ScreenTape Cat No 5067-5584) per manufacturer instructions. Average fragment size was 406 bp. Paired-end sequencing was performed on NovaSeq 6000 SP (2 x 100 paired end chemistry).

### Tumor Antigen Prediction and Computational Matrixing

Comprehensive tumor antigen discovery was performed using the LENS (Landscape of Effective Tumor antigens Software) pipeline version 1.7. Paired tumor and normal whole-exome sequencing (WES) along with tumor RNA-sequencing data were analyzed to identify tumor antigens arising from somatic mutations, cancer testis antigen, gene fusions, endogenous retroviruses, and viral sequences. Quality-controlled sequencing reads were aligned to the mouse reference genome (GRCm38) using BWA-MEM2 v2.1.1 for DNA and STAR v2.7.3a for RNA. Somatic variants were called using an ensemble approach combining Strelka2 v2.9.7, Mutect2 (GATK v4.6.1.0), and VarScan2 (v2.4.6), with variants retained if detected by any caller (union strategy). Germline variants were identified using DeepVariant v1.8.0. All variants were normalized, filtered using BCFtools v1.21, and phased with WhatsHap v2.4 to maintain haplotype information. The GENCODE vM28 gene annotation GTF was augmented with mouse-specific endogenous viral elements from gEVE. The resulting GTF file was using for both canonical transcript and ERV quantification using Salmon v1.10.3. Gene fusions were detected with STAR-Fusion v1.8.1b. Copy number alterations and tumor purity were estimated using CNVkit v0.9.9 and Sequenza v3.0.0, respectively. Tumor-associated and tumor-specific antigens were derived from multiple sources: somatic SNVs and InDels in expressed genes, cancer-testis antigens, endogenous retroviruses, viral peptides, and gene fusion breakpoints. Peptides of 8-11 amino acids were evaluated for MHC class I binding using NetMHCpan, MHCflurry, and NetMHCstabpan. All predicted tumor antigens were filtered against the reference proteome to retain only tumor-specific sequences and annotated with RNA-seq read support. Peptides were then prioritized for manual testing based upon their binding affinity, stability, and expression values. 145 peptides were selected based on their shared prediction in three biological replicates. Antigens were pooled for testing and deconvolution of results using the ACE software (56). Parameters were set where each peptide was represented in three distinct pools and each pool contained 4 to 5 peptides for a total of 90 pools.

### Logistic Regression Modeling

To identify peptide features associated with antigen immunogenicity, we performed univariable and multivariable logistic regression using Firth’s penalized likelihood approach. This was done to reduce bias from small sample size. Immunogenicity was defined as a binary outcome based on measurable T cell response via ELISpot positivity. T cell responses were considered immunogenic using RunDFR v1.13 through FredHutch (39, 40). Responses that met the threshold for significance for DFR(2x) were then considered immunogenic **(Supplemental File 1)**.

Peptide features of MHC binding affinity, RNA expression, peptide–MHC stability, and MHCflurry processing and presentation scores were evaluated in univariable models and multivariable models (39). Model performance was assessed using receiver operating characteristic curve analysis and area under the curve. All analyses were conducted in R statistical software using the logistf package.

### Pooled ELISpot Design and Deconvolution

In order to determine the immunogenic profiles of a large number of candidate peptides, we constructed a pooled ELISpot assay design using the ACE Configurator for ELISpot graphical user interface (ACE) (56). ACE, like other matrix design software (57) uses heuristic approaches to pool multiple peptides into wells where the overlap between pairs of peptides is minimized across replicates. This setup allows more efficient identification of immunogenic peptides by mapping pools that showed significant IFN-γ release to peptide memberships. ACE additionally uses deep learning to further optimize the assignment of peptides to pools by taking a sequence-aware approach. We used the ACE default parameters for the sequence thresholding.

ELISpot peptide pooling was done using ACE. Peptide pools of 4-5 antigens were generated to a total number of pools of 90. Peptide pooled ELISpots results were uploaded and deconvolved by ACE using the expectation maximization algorithm to nominate confident and candidate immunogenic peptides. Candidate and confident peptides were tested in single peptide ELISpost to confirm their immunogenicity.

### Peptide Ordering and Synthesis

A custom peptide library of 145 predicted antigenic peptides was ordered from Biosynth on a 2-umol scale. Antigens were then resuspended in DMSO, supplemented with molecular grade water into a 96-well plate of 2-mL tubes. Peptides were then further diluted in complete RPMI-1640 containing 10% FBS and 1x penicillin/streptomycin to achieve a final concentration in ELISpot of 1 nmol of each antigen per well.

### Tissue Collection and Processing

To enrich for CD8^+^ T cells for ELISpot testing, spleen and tumor draining lymph node (TDLN) samples were taken and dissociated. Spleens were dissociated using gentleMACS C tubes (Miltenyi Biotec, Cat No 130-093-237) on protocol spleen dissociation program 1 two times in 5-10 mL of complete RPMI-1640 (Gibco, Cat No 11875-093), 10% FBS, 1x penicillin/streptomycin, 1x L-glutamine, 55 uM 2-mercaptoethanol, 1x nonessential amino acids, and 1 mM sodium pyruvate. TDLN were dissociated using frosted glass slides in 5 mL of complete RPMI-1640 as stated above. Treatment groups were pooled together with respective TDLN and passed through a 70-um filter. Cells were pelleted at 1200 rpm for 5 min at 4 °C and resuspended in 1-5 mL of ACK Buffer (Gibco, Cat No A10492-01) and incubated for 2 min. 20 mL of complete RPMI-1640 was added to cells and pelleted as stated above. Cells were resuspended in 10 mL and run through 40-um filter. The filter was washed with 5 mL. Cells were pelleted and resuspended in 30-35 mL of media and counted using Thermofisher Countess II (diluted 1:1 cells to trypan blue). A kit was used for CD8^+^ T cell enrichment (Miltenyi Biotec, Cat No 130-104-075) via magnetic column, with calculations and procedures done according to the manufacturer instructions. Enriched CD8^+^ T cells were then resuspended in 1-20 mL of complete RPMI-1640 media and counted as stated above. Cells were resuspended to achieve desired cell count for ELISpot testing, and left over cells were frozen down in FBS with 10% DMSO.

### Matrixed ELISpot Set Up

ELISpot plates were coated with anti-IFN-γ antibody according to manufacturer instructions (BD, Cat No 551083) and blocked using complete RPMI-1640 for 2-3 hr on the day of ELISpot setup. Matrixed ELISpots were done in pooled animal studies. Splenocytes from tumor bearing animals 10 days post-tumor induction were processed as stated above and enriched for CD8^+^ T cells. DC2.4 cells were plated in ELISpots as antigen presenting cells at a 35:1 effector to target ratio. CD8^+^ T cells were plated at 100,000 cells per well. Excess CD8^+^ T cells were washed in 1x DPBS (Gibco, 14190-144) twice, frozen in FBS containing 10% DMSO, and stored in liquid nitrogen for tumor antigen validation ELISpot studies. Peptide pools contained 1 nmol of each antigen per pool. Each plate contained six technical replicates of negative controls (DMSO), three technical replicates of PHA (Thermo Scientific, R30852701) at 1x positive controls, and three technical replicates of OTI splenocytes with DC2.4s at experimental cell concentrations with 10 ug/mL of SIINFEKL (CPC Scientific, Cat No MISC-012A) as another control. Plates were incubated for two days, processed according to manufacturer’s instructions, and imaged on AID Gmbh plate reader HW ELR07 using software version 7.0 build 171453.

### Individual Peptide ELISpot Set up

ELISpot plates were coated and blocked as stated above in the matrixed ELISpot set up. Frozen cells from matrixed ELISpot studies were thawed in water bath at 37 °C and resuspended in warm complete RPMI-1640. CD8^+^ T cells were plated from 50,000-150,000 cells/well and DC2.4 cells were added as antigen presenting cells at 35:1 effector to target ratio. In single animal studies, freshly-enriched CD8^+^ T cells were used with the same parameters. ELISpot plates were incubated for two days with controls, processed, and imaged as above.

### Immunotherapy Administration

Fourteen days post-tumor cell injection, mice were given biweekly treatment of either anti-PD-1 (Bioxcell, clone RMP1-14, Cat No BE0146), isotype control (Bioxcell, clone 2A3, Cat No BE0089), or 1x PBS intraperitoneally at 10 mg/kg for two weeks. Seven days post-final treatment, enriched CD8^+^ T cells were pooled per treatment and assessed for reactivity against the 145 predicted antigenic peptides via matrixed ELIspot and individual peptide ELISpot as stated above. This experimental design was repeated for anti-CTLA4 (Bioxcell, clone 9D9, Cat No BE0164), isotype (Bioxcell, clone MPC-11, Cat No. BE0086), and 1x PBS at 10 mg/kg.

#### Protocol 1: Jaeger Lab Immunoprecipitation and Mass Spectrometry

##### Cell Culture of BBN963

BBN963 stocks were thawed and cultured in 150-mm plates and expanded to achieve 11 plates at 80% confluence. One plate was trypsinized with 1x 0.05% Trypsin-EDTA (Gibco, Cat No 25300-054) and counted to determine approximate cell count. The rest of the plates were scraped with 2 mL 1x DPBS (Gibco, Cat No 14190-144) to retrieve residual cells and added to suspensions. Each suspension was washed 2x with 1x DPBS and spun at 1280 rpm for 10 min at 4 °C. Cells were placed on ice and resuspended in 1 mL of 1x DPBS to achieve 100 x 10^6^ cells/mL. 1 mL of cell suspension was then put into a cryovial and spun down to remove excess supernatant, placed in dry ice for snap freezing, and put into LN_2_ storage for immunopeptidomics. 70 x 10^6^ BBN963 cells were treated with 20 ng/mL of murine IFN-γ (PeproTech, Cat No 315-05-100UG) for 48 hr to upregulate MHC-I and snap frozen as stated above for immunopeptidomics studies.

##### Antibody Conjugation for MHC-I Immunoprecipitation

For antibody conjugation to Protein-A Sepharose 4B beads (Thermo Fischer Scientific, Cat No 101042), 1-2 mL of 50% bead slurry was added to a microcentrifuge tube and spun for 2 min (100 rcf at 4 °C, same speed each following spin). Beads were washed with 1X PBS (Alkali Scientific, Cat No PB500) and spun for 2 min. 10 mg of anti-H2-Kb antibody (BioXcell, clone Y3, Cat No BE0172) or 10 mg of anti-H2-Db antibody (BioXcell, clone 28-14-8S, Cat No BE0451) per 1 mL of bead resin in PBS was added and the sample was rotated at room temperature for 1 hr. After rotating, the beads were spun and washed once with 1X PBS and 0.2 M sodium borate pH 9.0 (Thermo Fisher Scientific, Cat No J63637.AK). Antibody-bound beads were resuspended in 40 mM dimethyl pimelimidate dihydrochloride (DMP) (Sigma-Aldrich, Cat No D8388) in 0.2 M sodium borate and incubated with rotation for 30 min at room temperature. Beads were spun and resuspended again in a new round of 40 mM DMP. After incubation, the bead pellet was quenched with 0.2 M ethanolamine (Sigma-Aldrich, Cat No 398136) in 0.2 M sodium borate and rotated for 2 hr at room temperature. Beads were spun and washed with the following in order: 100 mM Tris pH 8.0 (Invitrogen, Cat No AM9856), 100 mM glycine in HCl pH 2.5-4.0 (Thermo Fischer Scientific, Cat No, Sigma-Aldrich, 320331), 1X PBS. After washing, beads were resuspended to create a 1:1 bead to PBS slurry and were stored in 4 °C.

##### Extraction of MHC-I Complexes

For *in vitro* samples, cells were treated with IFN-γ (20 ng/ml) for 48 hr prior to collection. Flash frozen cell pellets were resuspended in 1.75 mL of MHC Extraction Buffer (MEB), 20 mM Tris pH 8, 1 mM EDTA, 100 mM NaCl, 1% Triton X 100, 60 mM Octyl beta-D-glucopyranoside (OGP), 6 mM MgCl_2_, 0.2 mM Iodoacetamide, 1X HALT protease inhibitors, 250 U/µl Benzonase (Sigma Aldrich) and rotated for 30 min at 4°C. Debris was pelleted at 14,000 x *g* and supernatant moved to fresh protein Lo-bind tube. For each sample, a mixture of 50 μL H2-K^b^ and 50 μL H2-D^b^ bead slurry (Fisher Scientific) was equilibrated using MEB buffer, and was then added to the cleared lysates, and rotated at 4 °C for 3 hr. Following this, beads were then washed twice with MEB buffer, twice with 1x PBS, and then twice with 10 mM Tris pH 8. To elute complexes from beads, samples were incubated with 200 μL 1% Trifluoroacetic acid (TFA) at room temperature for 10 min. Beads were then centrifuged at 16,000 x *g* for 10 min and the resulting supernatants were then desalted with C18 spin tips. T3 200 C18 Affinisep tips (Affinisep) were prepared for peptide clean up by first applying acetonitrile (©) + 0.1% Formic Acid (FA) to tip matrix and centrifuging through tips. 80% ACN 0.1% FA mixture was then added and the sample centrifuged. Water with 0.1% FA was then added and the sample centrifuged. Peptide containing supernatants were then added to the tip matrix and centrifuged. Tips were then washed by adding 3% ACN, 0.1% FA and samples were centrifuged. Finally, peptides were eluted from the tips twice with 7.5 μL 50% ACN, 0.1% FA and centrifugation directly into autosampler vials. Solvents were then removed via Speedvac and peptides reconstituted in 2% ACN, 0.1 mM Peptide Retention Time Calibration, and in 0.1% FA prior to analysis by LC-MS/MS.

##### Liquid Chromatography-Tandem Mass Spectrometry

MHC peptides were analyzed using a Thermo Orbitrap Ascend Tribrid mass spectrometer (Thermo Scientific) coupled with a Vanquish Neo UHPLC system (Thermo Scientific) and equipped with a FAIMS Pro Duo. Samples were resuspended in 2% ACN + 0.1% FA and directly loaded onto a precolumn (2 cm x μm ID packed with C18 reversed-phase resin, 5 μm, 100 Å) in solvent A (2% ACN, 0.1% FA). Trapped peptides were eluted onto the analytical capillary chromatography column with an integrated electrospray tip (C18, 75 μm ID × 25 cm, 2 μm, 100 Å, Thermo Scientific). Peptides were eluted over 105 min with the following: gradient 2–3% solvent B (90% ACN, 0.1% FA) for 10 min, 3–5% solvent B for 2 min, 5–23% solvent B for 63 min, 23-40% solvent B for 22 min, 40-90% solvent B for 3 min, hold for 5 min and followed by column equilibration and wash.

Data dependent acquisition was performed in positive ion mode at a spray voltage of 2.0 kV and heated capillary temperature, 275 °C. Full scan mass spectra (300–1,000 *m/z*, 120,00 resolution for MHC-I and 300-1,200 m/z) were detected in the Orbitrap analyzer after accumulation of 2.5 × 10^6^ ions (normalized automatic gain control (AGC) target of 250%) and 250 ms maximum injection time (IT). For every full scan, MS^2^ spectra were collected during a 1.5 s cycle time for FAIMS compensation voltages at -45 and -65. Precursor ions were isolated with a 2.0 m/z isolation window, and accumulation time was set to auto with 250 ms maximum accumulation time. Ions were fragmented with higher energy disassociation collision (HCD) with 30% collision energy (CI) at a resolution of 30,000. Charge states <2 and >4 were excluded, and dynamic exclusion was set to 15 s after *n* = 1 observation. For acquisitions with an inclusion list, the precursor list included all peptides of interest with charge states 2-4 and the mass tolerance for inclusion was set to 15 ppm.

##### LC-MS/MS Database Searching

DDA MS searching was performed with FragPipe (v.21.0) using the HLA-workflow against a custom database containing the Gencode v36 murine proteome and somatic mutations previously defined above. Group-specific FDR was set to 1%. The search was completed with non-specific protein digestion, methionine oxidation, cysteine carbamidomethylation, and protein N-term acetylation set as variable modifications along with peptide length at 8-11 amino acids. Label free quantification was performed using IonQuant using precursor abundance. Match between runs (MBR) was set to true with an FDR of 1%. Search outputs were subsequently analyzed and visualized in R or Skyline. Manual spectrum inspection was performed in PDV v2.2.

#### Protocol 2: Biognosys Immunoprecipitation and Mass Spectrometry

##### Immunopeptidomics Sample Preparation

Snap frozen BBN963 cell pellets were submitted to Biognosys for murine MHC Class I immunopeptidome profiling. These samples and a processing quality control sample (frozen mouse lung tissue) were lysed using Biognosys’ Lysis Buffer for 1 hr at 4 °C. After removal of cell debris, the lysates were incubated with bead-coupled antibodies (Clone M1/42.3.9.8) to capture MHC class I complexes with associated peptides. After immunoprecipitation, bound protein complexes were washed and peptides eluted using Biognosys’ optimized immunopeptidomics protocol. Eluted immunopeptides were filtered using a 10 kDa MWCO plate (Millipore). Clean up for mass spectrometry was carried out using an Oasis HLB µElution Plate (30 µm, Waters) according to the manufacturer’s instructions. Immunopeptides were dried down using a SpeedVac system and resuspended in 10 µL resuspension solvent (1% acetonitrile/0.1% formic acid (FA) in water supplemented with 0.015% DDM). The spectral library pool samples were pooled and fractionated into 5 fractions using high pH reversed phase fractionation (HPRP) for spectral library generation. Peptide concentrations in mass spectrometry ready samples were measured using a Fluorometric Peptide Assay (Pierce, Thermo Scientific).

##### DDA Mass Spectrometry Acquisition

For DDA LC-MS/MS measurements, 6 µL of immunopeptides per sample were injected on a custom Aurora-3 series Frontier (IonOpticks) reversed phase column on a Thermo Scientific NeoVanquish UHPLC nano-liquid chromatography system connected to a Thermo Scientific™ Orbitrap™ Exploris 480™ mass spectrometer equipped with a Nanospray Flex™ ion source and a FAIMS Pro™ ion mobility device (Thermo Scientific™). LC solvents were A: water with 0.1% FA; B: 80% acetonitrile, 0.1% FA in water. The nonlinear LC gradient was 1-44.3% solvent B in 120 min followed by a column washing step in 90% B for 8 min, and a final equilibration step of 1% B for 11 min at 64°C with a flow rate set to 350 nL/min. MS1 precursor scans were acquired between 3–0 - 800 m/z at 120,000 resolution with data-dependent MS2 scans at 60,000 resolution. MS2 scans were acquired in cycle time of 2.03 s per applied FAIMS compensation voltage. Only precursors with charge state 1-5 were isolated for dependent MS2 scans.

##### DIA Mass Spectrometry Acquisition

For DIA LC-MS/MS measurements, 150 ng of immunopeptides per sample were injected on a custom Aurora-3 series Frontier (IonOpticks) reversed phase column on a Thermo Scientific NeoVanquish UHPLC nano-liquid chromatography system connected to a Thermo Scientific Orbitrap Exploris 480 mass spectrometer equipped with a Nanospray Flex™ ion source and a FAIMS Pro ion mobility device (Thermo Scientific). LC solvents were A: water with 0.1% FA; B: 80% acetonitrile, 0.1% FA in water. The nonlinear LC gradient was 1-44.3% solvent B in 120 min followed by a column washing step in 90% B for 8 min, and a final equilibration step of 1% B for 11 min at 64 °C with a flow rate set to 350 nL/min. The FAIMS-DIA method consisted per applied compensation voltage of one full range MS1 scan and 8 DIA segments as adopted from Bruderer et al. (58) and Tognetti et al. (59).

##### Database Search of DIA and DDA LC-MS/MS Data and Hybrid Spectral Library Generation

The DIA mass spectrometric data (3 runs) and the shotgun mass spectrometric data (5 fractions) were searched separately. The two datasets were searched using Spectronaut (Biognosys, version 20.2). The false discovery rate on peptide level was set to 1% for each dataset. A mouse UniProt .fasta database (*Mus musculus*, UP000000589, 2025-07-01) and the custom databases were used as a search database. Non-specifically digested peptides (no trypsin-cleaved termini) with a length between 7 and 14 amino acids, with max. 2 variable modifications (N-term acetylation, oxidation (M), cysteinylati©(C), carbamidomethyl©on (C)) were included in the search. A hybrid spectral library was constructed from the two search archives, with the false discovery rate on peptide level set to 1%.

##### DIA Data Analysis

Raw DIA mass spectrometric data were analyzed using Spectronaut (Biognosys, version 20.2) with q-value filtering and no background signal integration or imputation. The hybrid spectral library generated in this study was used. Settings included peptide level false discovery rate control at 1%. The DIA measurements analyzed with Spectronaut were normalized using local regression normalization (60).

##### Data Analysis

Downstream data analysis including peptide length distribution and peptide motifs was performed in R. Distance in heat maps was calculated us42anhattanmanhattan” method, the clustering using “ward.D” for both axes. General plotting was done in R using ggplot2 package.

##### Single Cell GEM Generation and Sequencing Library Preparation

BBN963 Single cell barcoded cDNA was generated using the Chromium Next GEM Single Cell 5’ Kit v2 (10x Genomics). Sample suspensions were loaded into separate lanes of a Chromium Next GEM K Chip (10x Genomics) at a target cell recovery of 10,000 cells per sample. The cells were partitioned into individual gel beads-in-emulsion (GEMS) using the 10x Genomics Chromium Controller. RNA from encapsulated cells in gel beads coated with barcoded oligos were reverse transcribed using a Veriti 96-well thermal cycler (Applied Biosystems). Complementary DNA was thereby labeled with a cell barcode and unique molecular index (UMI). The GEMs were broken, and the unamplified cDNA was purified from the emulsion mixture using MyOne Silane Dynabeads (Invitrogen). Purified cDNA was amplified for 13 cycles, and subsequently purified with SPRIselect size selection beads (Beckman Coulter). cDNA concentration was determined on a Qubit 4 Fluorometer using the Qubit dsDNA high sensitivity kit (ThermoFisher). Size distribution of the cDNA was verified on the TapeStation 2200 with the High Sensitivity D5000 reagents (Agilent Technologies).

Fifteen nanograms of single-cell cDNA generated using the 10x Chromium Next GEM Single Cell 5’ Kit v2 was used in a cDNA amplification reaction with Kinnex capture primers, followed by Template-Switch-Oligo artifact removal. The cDNA was subsequently amplified with 16 Kinnex PCR reactions with Kinnex primers to generate DNA fragments containing orientation-specific Kinnex segmentation sequences. Then PCR-amplified Kinnex cDNA fragments were pooled equally and treated with Kinnex enzyme and then ligated to barcoded Kinnex terminal adapters to assemble cDNA segments into an array. After removal of incomplete array, the final Kinnex library was bound to SPRQ polymerase with SPRQ Polymerase Kit, then sequenced with SPRQ Sequencing Plate and Revio SMRT cell (25M) on the Revio instrument for 24 hr. Libraries for Illumina sequencing were generated with 50 ng of cDNA using the 10x Genomics 5’ GEX-X Gene Expression Library Construction protocol with no modifications (10x Genomics). Libraries were sequenced on an Illumina NovaSeq 6000 using 150 bp PE reads.

##### Long Read Single Cell Sequencing Data Analysis

Raw sequencing data were processed using the standard PacBio pipeline to generate circular consensus sequences (CCS) with ccs v8.2.0. HiFi reads were trimmed using pbtrim v1.1.0 and demultiplexed with lima v2.11.0 using the SYMMETRIC-ADAPTERS preset. Base modifications (6mA) were called using jasmine v2.3.0. Concatenated full-length transcript reads were segmented into individual transcripts using skera v1.3.0. The segmented reads were then processed through the single-cell IsoSeq pipeline, which included refinement, error correction, and deduplication using IsoSeq groupdedup v4.2.0. Finally, deduplicated transcripts were aligned to the mouse reference genome (GRCm39/mm39 with GENCODE vM28 annotations) using pbmm2 v1.16.0 with the ISOSEQ preset, and alignments were coordinate-sorted using SAMtools v1.19.1.

Seurat-compatible input files were generated using Pigeon. These files were then converted to gene-level counts using a custom script. The resulting matrix was filtered to the set of cancer-testis antigens generating immunogenic pMHCs. Separately, cells were also categorized for somatic variant presence using a custom script. Specifically, immunogenic somatic variants were lifted over from the standard GRCm38 reference to the GRCm39 reference. Next, a custom script was used to, for each cell, determine whether any of the cell-specific reads covering the genomic origin of the variant contained the alternate allele. Cells with at least one read supporting the variant were classified as containing the variant, or were classified as wildtype otherwise. Endogenous retroviruses generating immunogenic peptides were lifted over from GRCm38 coordinates to GRCm39 coordinates, aggregated into a custom GTF file, and then quantified using Pigeon. The resulting SNV, ERV, and CTA matrices were merged by cellular barcode. Each cell’s tumor antigen status was reduced to a binary representation by converting read support greater than 0 (expression read count for ERVs and CTAs; alternate allele read count for SNVs) to “True” and read support of zero to “False”.

A greedy selection algorithm was used to determine potential tumor cellular coverage by sets of pMHCs selected to maximize coverage. Tumor antigen networks were constructed by aggregating each cell’s collection of tumor antigen statuses into a binary vector. Binary vectors that were observed in 50 or more cells were used as nodes within the network plot with edges connecting nodes with a Hamming distance of 1.

## Supporting information

DFR2x Analysis

Supplemental Figure 1 and 2 with Table 1 and Table 2

## Acknowledgements

University of North Carolina Immune Monitoring and Genomics Facility (IMGF)

University Cancer Research Fund (UCRF)

UNC Cancer Center Core Support Grant #P30CA016086

Institute for Genome Sciences, University of Maryland School of Medicine

Jamie Leandro Foundation

NIH/NCI R01 CA241810 (BGV, WYK)

NIH/NCI R37 CA247676 (BGV)

Visualization code and figure design for Figure 3C and Supplemental Figure 2 were developed with the assistance of Claude Sonnet 4.6, a large language model (Anthropic, 2025); all scientific interpretations, analytical decisions, and conclusions are solely those of the authors.

OpenAI’s GPT-5.5 Thinking was used to assist with copyediting and improving the clarity and readability of selected manuscript text; all analyses, interpretations, scientific conclusions, and intellectual content are original work of the authors.

## Data Availability Statement

The sequencing data, LENS prediction results, raw ELISpot data, and Mass Spectrometry data generated and analyzed in this study are available from the corresponding authors upon request.

## Notes

### Competing Interest Statement

The authors have declared no competing interest.

### Summary of Updates

Authors have been updated. Figure 6 has been updated to reflect current immunogenic antigen calls based on the statistical testing criteria.

