## Supplemental Figure 1 and 2 with Table 1 and Table 2 for "Immunodominance Hierarchy of Endogenous BBN963 Bladder Cancer Antigens Remains Stable Under Anti-PD1 and Anti-CTLA4 Immunotherapy"

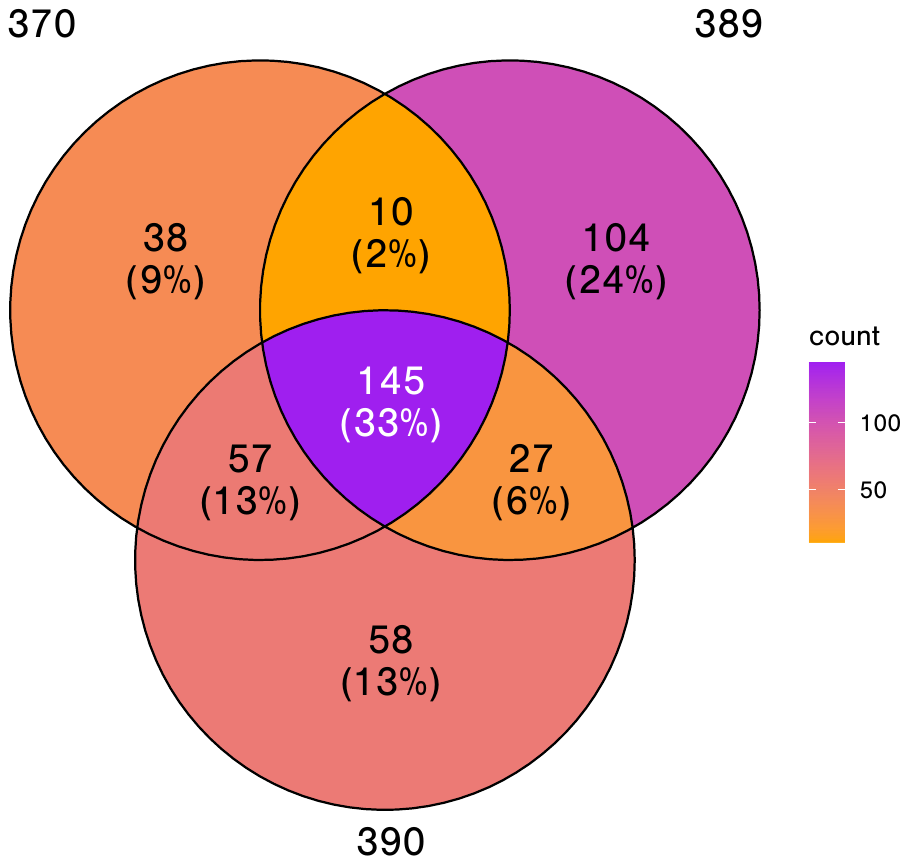

**Supplemental Figure 1. Shared LENS Predicted Tumor Antigen Peptides Among Three BBN963 Tumor Replicates.** Venn diagram contains number of antigens found in only one replicate, at the intersection of two replicates or in all three (denoted by white font).

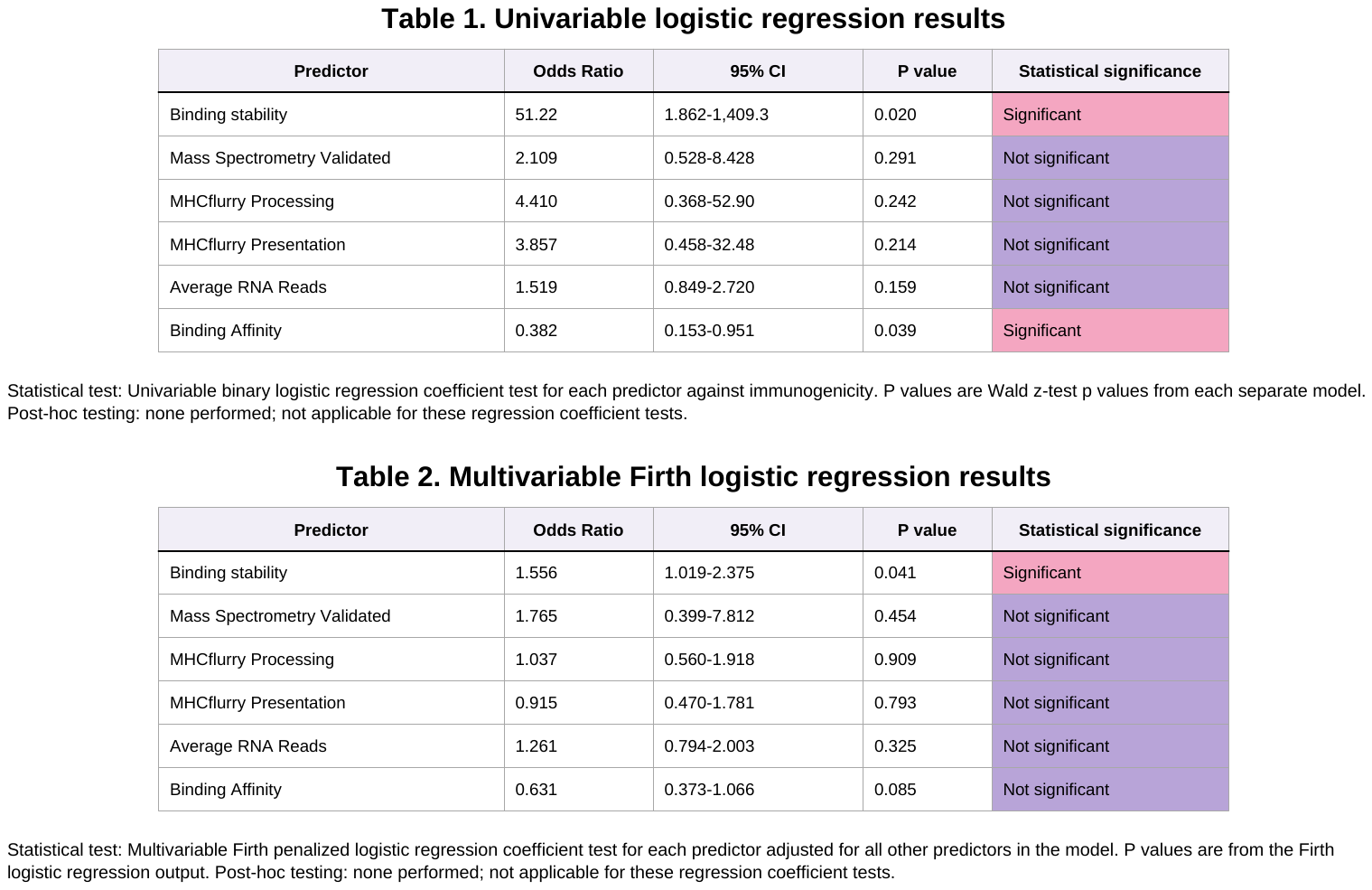

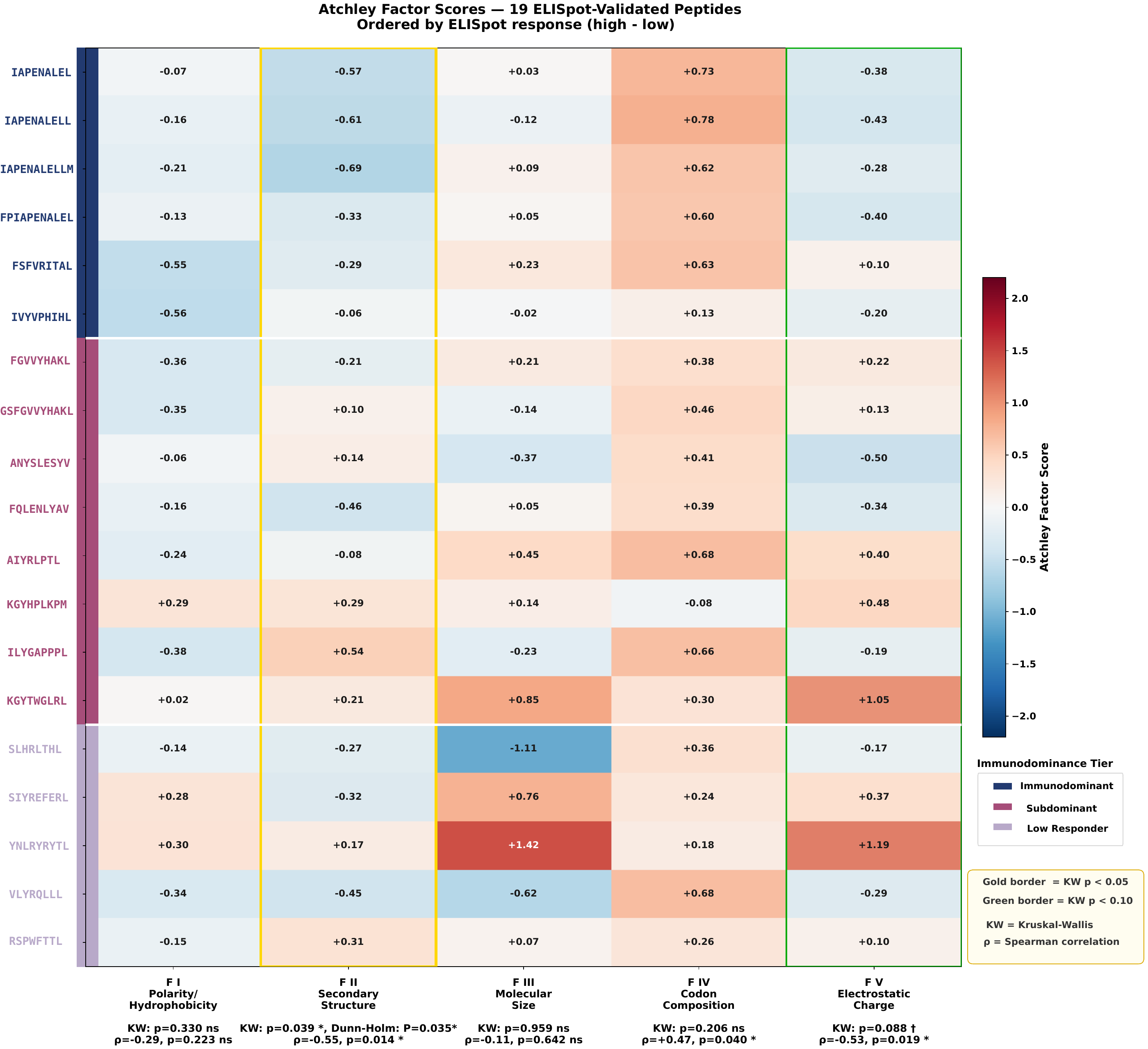

**Supplemental Figure 2. Heatmap displaying mean Atchley factor scores across five physiochemical axes for 19 validated tumor antigens**. Scores were computed by averaging per residue Atchley factor across all positions of each peptide. Gold column border indicates significant separation of immunodominance tiers by Kruskal-Wallis test with Dunn-Holm post hoc (P <0.05). Green border indicates trending results (P <0.10). Left color strip denotes immunodominance tier: navy (immunodominant, >500 SFU), berry (subdominant, 100–500 SFU), lavender (low responder, <100 SFU), relative to DMSO negative control (5 SFU/10⁶ cells)
